# A neuroendocrine feedback loop in *C. elegans* males integrates food detection and biological sex to modulate chemoreceptor expression and behavioral flexibility

**DOI:** 10.1101/752238

**Authors:** Leigh R. Wexler, Renee M. Miller, Douglas S. Portman

## Abstract

Dynamic integration of internal and external cues is essential for flexible, adaptive animal behavior. In *C. elegans*, biological sex and feeding state regulate expression of the food-associated chemoreceptor *odr-10*, contributing to plasticity in food detection and the decision between feeding and exploration. In adult hermaphrodites, *odr-10* expression is high; in well-fed adult males, *odr-10* expression is low, promoting exploratory mate-searching behavior. Food-deprivation transiently activates male *odr-10* expression, heightening food sensitivity and reducing food-leaving. Here, we identify a neuroendocrine feedback loop that sex-specifically regulates *odr-10* in response to food deprivation. In well-fed males, insulin-like (IIS) and TGFβ signaling repress *odr-10* expression. Upon food deprivation, *odr-10* is directly activated by DAF-16/FoxO, the canonical *C. elegans* IIS effector. The TGFβ ligand DAF-7 acts upstream of IIS, and, likely because of its sexually dimorphic expression in the nervous system, links feeding to *odr-10* only in males. Surprisingly, these responses to food deprivation are not triggered by internal metabolic cues, but rather by the loss of sensory signals from food. In the presence of inedible food, males become metabolically starved but express levels of *odr-10* and *daf-7* comparable to those of well-fed males. Further, exposing food-deprived males to inedible food is sufficient to restore low *odr-10* expression. Food signals are detected by a small number of sensory neurons whose activity non-autonomously regulates *daf-7* expression, IIS, and *odr-10*. Thus, adult *C. elegans* males employ a neuroendocrine feedback loop that integrates food detection and genetic sex to dynamically modulate chemoreceptor expression and sensory behavior.

## INTRODUCTION

Innate drives—like those to feed, reproduce, and avoid predators—provide a flexible and modular framework for the control of animal behavior. Internal and external signals continuously interact with such drives, allowing behaviors to be activated or suppressed according to an animal’s history, physiology, and environment. In humans, disruption of these mechanisms is associated with a variety of neuropsychiatric conditions, including eating disorders, depression, and obsessive-compulsive disorder. Though even the simplest nervous systems employ flexible, innate drives, much remains unknown about how they are regulated by internal and external conditions, how these conditions are represented internally, and how drive states influence behavioral choice.

One mechanism by which internal drives and states influence behavior is through modulation of sensory perception. Feeding state is a primary source of such regulation: for example, hunger and satiety modulate chemosensory function in *Drosophila* [1–4], *C. elegans* [5–12], and mice [13, 14], as well as mechanosensation in the leech [15]. In this way, hunger focuses an animal’s attention on food-related appetitive stimuli and attenuates the perception of cues that might otherwise inhibit feeding [16, 17]. In most of these cases, feeding state information is thought to originate from internally derived metabolic cues, with the role of acute sensory input in influencing internal state being less well appreciated.

Other dimensions of internal state, like biological sex and reproductive status, also influence sensory function [18]. In *C. elegans*, for example, biological sex modulates sensory responses to pheromones and food [12, 19–23]. In *Drosophila*, chemoreceptor expression differs by sex [24–26] and chemosensory processing is modulated by female reproductive state [27]. In mice, biological sex shapes sex differences in pheromone detection [28] and hormonal signals of female reproductive status can act directly on sensory neurons to influence their sensitivity to pheromonal and olfactory cues [29–31]. However, how biological sex and reproductive state intersect with feeding status and other dimensions of internal state to adaptively modulate behavior is largely unexplored.

State-dependent food-leaving behavior in *C. elegans* provides an ideal context for investigating the mechanisms by which multiple internal and external cues are integrated to balance drives and influence behavioral choice [32]. Under most circumstances, individual worms are efficiently retained by a small spot of bacterial food. However, well-fed adult males tend to leave a food source long before it is depleted, abandoning it in search of a suitable mate [23]. Multiple mechanisms are important for the prioritization and execution of this exploratory behavior, including input from male-specific sensory neurons that may tonically signal the absence of potential mates [33] and peptidergic signaling through the PDFR-1 receptor [34, 35]. Additionally, adult males diminish their attraction to some food-related cues compared to adult hermaphrodites and larval males, thereby promoting exploration over feeding [12, 21]. This occurs in part through repression of *odr-10*, the chemoreceptor for the food-associated odorant diacetyl [12, 36]. Downregulated *odr-10* expression in adult males is influenced by the genetic sex of the AWA neurons, the chemosensory neuron pair that expresses this receptor [12], as well as a temporal cue signaling the transition to adulthood, controlled by the heterochronic genes *lep-2* and *lep-5* [37].

Food-leaving behavior in adult males is also subject to homeostatic regulation, balancing the drive to mate with the drive to feed. If males are deprived of food for several hours and then placed on a food source, food-leaving behavior is transiently suppressed, returning to baseline levels only after several hours of re-feeding [23]. Food-deprivation transiently activates *odr-10* expression in males and increases *odr-10*-dependent food attraction, indicating that sensory plasticity is a component of this behavioral switch [12].

To learn more about state-dependent regulation of chemoreceptor expression and sensory behavior, we investigated the mechanisms that activate *odr-10* expression upon food deprivation in *C. elegans* adult males. In doing so, we identified a series of regulatory interactions that couple feeding to low *odr-10* expression, together constituting a neuroendocrine feedback loop involving the TGFβ-superfamily signal DAF-7, the Insulin-IGF-1-like signaling (IIS) receptor DAF-2, and its primary effector, the FoxO transcription factor DAF-16. Further, we find that *odr-10* is a direct target of DAF-16 in the AWA neurons. This mechanism is largely male-specific, likely as a consequence of the recently described male-specific expression of *daf-7* in the ASJ neurons [38]. *odr-10* expression is also increased in response to food-deprivation in hermaphrodites, but this increase is primarily mediated by distinct mechanisms. Surprisingly, the signal that initiates this loop in males is not an internal metabolic cue: starvation has little effect on *odr-10* if it occurs in the presence of food stimuli, but blocking the perception of food cues is sufficient to activate *odr-10* expression even in well-fed males. Together, these findings identify a sex-specific neuroendocrine feedback mechanism that couples sensory perception to chemoreceptor expression, thereby contributing to the homeostatic balance of behavioral drives.

## RESULTS

### *Insulin signaling regulates* odr-10 *in response to feeding state in males*

In adult *C. elegans* males, expression of *odr-10* in the AWA neurons is transiently activated by 12-18 h of food deprivation, returning to low baseline after several hours of re-feeding [12] (Figs. 1A, B). To identify mechanisms by which food availability regulates *odr-10*, we examined *odr-10* expression in males carrying mutations in regulatory pathways previously associated with feeding state. We first considered the insulin/IGF-1 signaling (IIS) pathway, since the activity of *daf-2*, the receptor for the *C. elegans* IIS pathway, promotes food-leaving behavior in well-fed males [23]. Using the translational *ODR-10::GFP* fosmid reporter *fsEx295* [12], we found that well-fed *daf-2* mutant males exhibited high *odr-10* expression, comparable to that seen in food-deprived wild-type males (Fig. 1C). This upregulation was eliminated in *daf-16; daf-2* double mutants, indicating that the activation of *odr-10* by reduced insulin signaling depends on *daf-16*, the FoxO transcription factor that is the canonical effector of IIS in *C. elegans* (Fig. 1C). Further, we found that loss of *daf-2* significantly increased males’ attraction to diacetyl, the ligand for *odr-10* (Fig. 1D).

**Figure 1.**
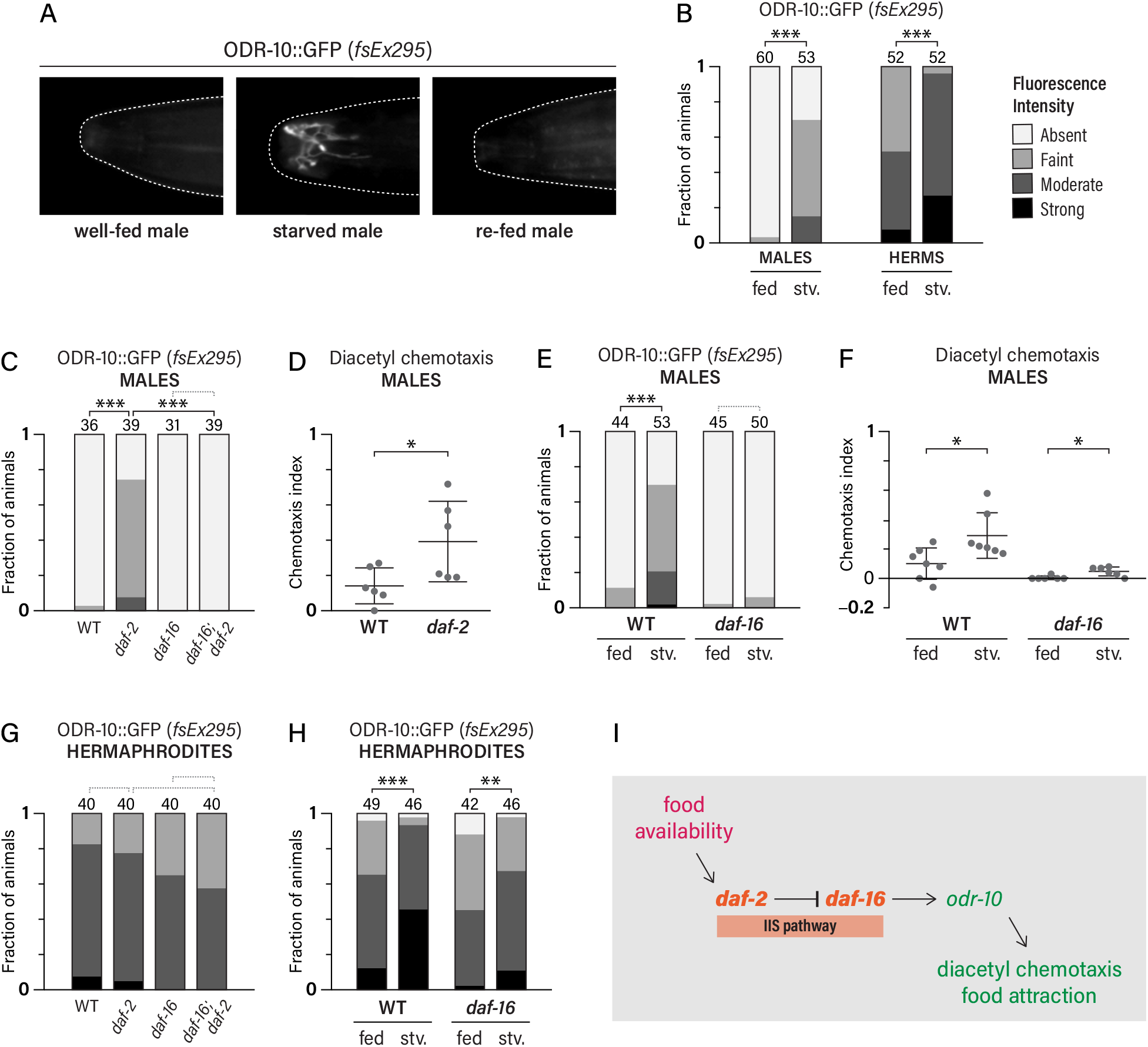
Insulin-like signaling regulates *odr-10* expression males in response to feeding state. (A) Representative images of ODR-10::GFP fluorescence in a well fed male, a food-deprived (“starved”) male, and a male re-fed after food deprivation. (B) ODR-10::GFP expression in WT fed and starved (stv) males and hermaphrodites, scored qualitatively on a four-point scale (see Methods). (C) ODR-10::GFP expression in male insulin-signaling mutants. (D) Chemotaxis to diacetyl (1:1000) in WT and *daf-2* males. (E) ODR-10::GFP expression in fed and starved WT and *daf-16* males. (F) Chemotaxis to diacetyl (1:1000) in WT and *daf-16* males either well-fed or starved for 12 hours. (G) ODR-10::GFP expression in hermaphrodite insulin-signaling mutants. (H) ODR-10::GFP expression in fed and starved WT and *daf-16* hermaphrodites. (I) Pathway for the regulation of *odr-10* by food availability and IIS. *0.01 < *p* < 0.05; **0.001 < *p* < 0.01; \*\*\**p* < 0.001. Dotted gray brackets indicate *p* > 0.05.

To ask whether the upregulation of *odr-10* by food deprivation could be caused by a change in IIS, we examined *ODR-10::GFP* expression in food-deprived *daf-16* males. Unlike wild-type, *daf-16* males exhibited no change in reporter expression upon food deprivation (Fig. 1E). Similarly, starvation elicited a large increase in diacetyl attraction in wild-type males but had only a very minor effect in *daf-16* mutants (Fig. 1F). Together, these findings indicate that food deprivation upregulates *odr-10* through decreased IIS and increased *daf-16* activity. However, they do not rule out the possibility that *daf-16* is simply a permissive activator, necessary for *odr-10* expression under all circumstances. To test this, we examined larval males. As shown previously, L3 males exhibit high *odr-10* expression, comparable to that of L3 hermaphrodites, with low *odr-10* expression in males emerging only upon the transition to adulthood [12, 37]. We found that larval *odr-10* expression was largely unaffected in *daf-16* mutant males (Fig. S1A), ruling out a general requirement of *daf-16* for all *odr-10* expression. This indicates that an increase in *daf-16* activity likely underlies the activation of *odr-10* in food-deprived adult males (Fig. 1I).

### *The response of* odr-10 *to food deprivation differs by sex*

Our previous work indicated that the induction of *odr-10* expression by food deprivation occurred only in males [12]. However, by qRT-PCR, we observed a significant increase in *odr-10* mRNA upon food deprivation in hermaphrodites (Fig. S1B). Furthermore, a recent report found that *odr-10* expression was increased in *daf-2* mutant hermaphrodites [39]. Therefore, we considered the possibility that *kyIs53*, the *ODR-10::GFP* reporter used previously, might be prone to ceiling effects that could prevent the detection of increased fluorescence in food-deprived hermaphrodites. When we examined hermaphrodites carrying the *ODR-10::GFP* fosmid reporter *fsEx295*, whose baseline expression level is lower than that of *kyIs53*, we found that food deprivation did indeed elicit a significant increase in GFP intensity (Fig. 1B). Thus, *odr-10* expression is sensitive to food availability in both sexes, providing a homeostatic mechanism by which the absence of food could increase in an animal’s sensitivity to food odor.

Despite this qualitatively similar response to food deprivation, however, we observed sex differences in the underlying mechanisms. Unlike males, *daf-2* mutant hermaphrodites exhibited no apparent change in *ODR-10::GFP* intensity (Fig. 1G). Consistent with this, we found that *daf-16* was not required for the starvation-induced increase of *odr-10* expression in hermaphrodites (Fig. 1H). The apparent discrepancy between our findings and the qRT-PCR results of Hahm *et al*. [39] might indicate an important role for post-transcriptional regulation of *odr-10* in hermaphrodites; further studies are needed to evaluate this and other possibilities. Nevertheless, these findings demonstrate that IIS is not the primary mediator of *odr-10* activation by food deprivation in hermaphrodites and that the sexes differ in the nature of the mechanisms that link feeding state to *odr-10* expression.

### odr-10 *is likely a direct target of* daf-16

To better understand how IIS regulates *odr-10*, we asked whether *daf-16* acts cell-autonomously. We found that expression of *daf-16f* cDNA under the control of the AWA-expressed promoter *Pgpa-4Δ6* [12] was sufficient to restore *odr-10* expression in starved *daf-16* males (Fig. 2A). Consistent with this, we detected *DAF-16::GFP* in AWA nuclei in *daf-2* mutant males (Fig. 2B). In contrast, intestine-specific expression of *daf-16* had no effect on *odr-10* (Fig. S2A, B).

**Figure 2.**
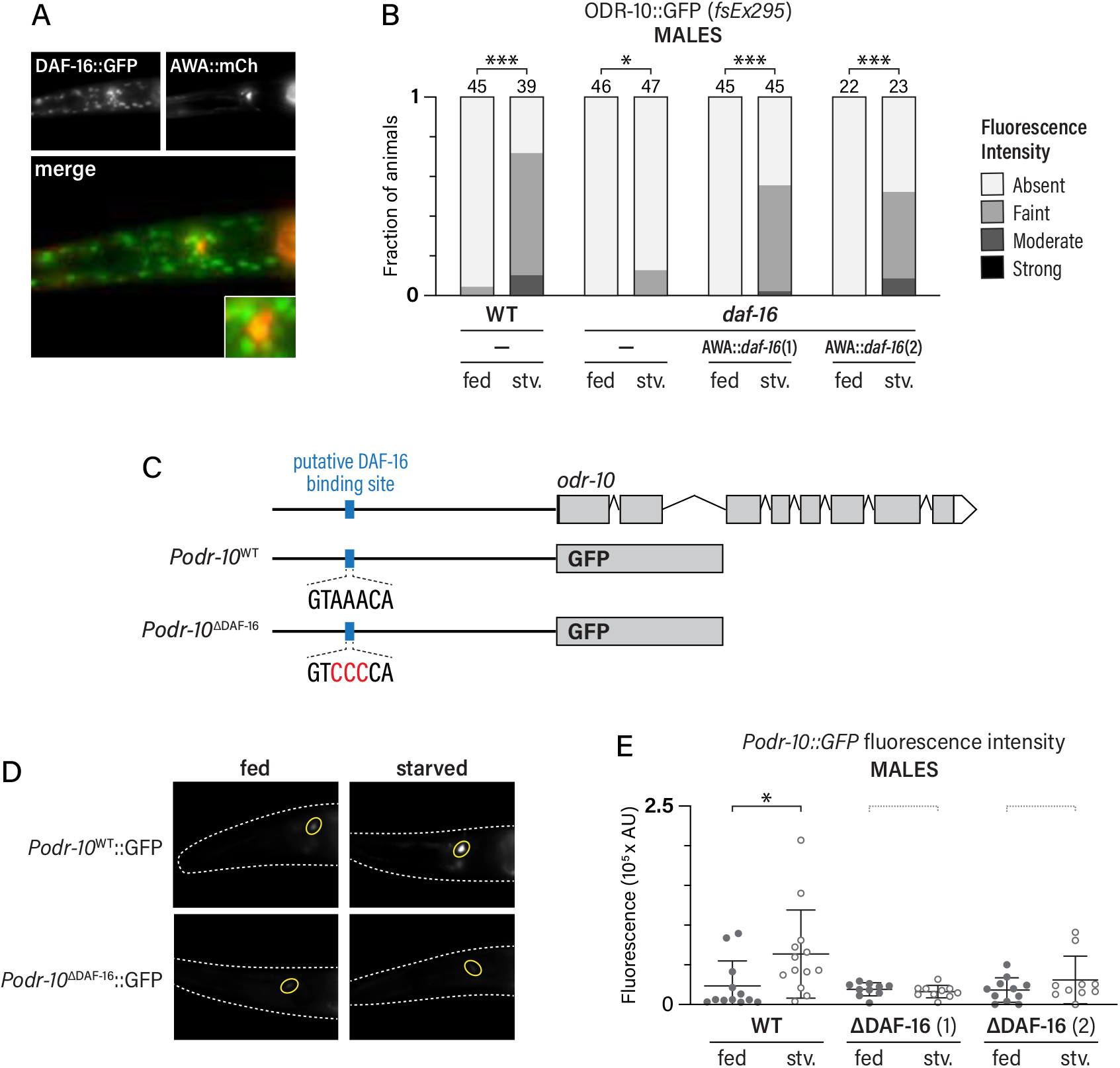
DAF-16 acts cell-autonomously, and likely directly, to regulate *odr-10*. (A) DAF-16::GFP and AWA::mCherry expression. Upper panels show individual channels; lower panel shows a merged, pseudocolored image with a higher-magnification inset. To bring about nuclear localization of DAF-16, this strain carried a *daf-2* mutation and was grown at 25°C. (B) ODR-10::GFP expression in WT and *daf-16* fed and starved (stv) males, with or without (–) *AWA::daf-16f* transgenes. Results are shown for two independent lines. (C) Diagram depicting the *odr-10* genomic locus (upper) and *odr-10* transcriptional reporters (middle and lower) with wild-type and mutant versions of the putative DAF-16 binding site. (D) Representative images of *Podr-10^WT^::GFP* and *Podr-10^ΔDAF-16^::GFP* in fed and starved males. Yellow ovals indicate the location of AWA. (E) Quantification of GFP fluorescence in *Podr-10^WT^::GFP* and *Podr-10^ΔDAF-16^::GFP* in fed and starved males. For the mutant reporter, results using two independent lines are shown. *0.01 < *p* < 0.05; **0.001 < *p* < 0.01; \*\*\**p* < 0.001. Dotted gray brackets indicate *p* > 0.05.

These results raised the possibility that *odr-10* might be a direct transcriptional target of DAF-16. Supporting this, a putative DAF-16 binding element was recently identified 809 bp upstream of the *odr-10* start codon [39]. To ask whether DAF-16 might act through this site, we generated two *Podr-10::GFP* reporters, one containing roughly 1 Kb of the wild-type *odr-10* promoter (*Podr-10*^WT^) and one in which this element was disrupted by changing three conserved nucleotides (*Podr-10*^ΔDAF-16^) (Fig. 2C). We observed a significant induction of *Podr-10*^WT^::GFP expression upon food deprivation, but detected no change in *Podr-10*^ΔDAF-16^::GFP (Fig. 2D, E). These results show that the induction of *odr-10* by food deprivation in males is mediated at least in part at the transcriptional level. Moreover, they show that DAF-16 functions in AWA and that *odr-10* is likely to be a direct target of the *C. elegans* IIS pathway.

### *TGFβ signaling acts upstream of IIS to regulate* odr-10 *in males*

We also considered a role for the BMP/TGFβ-family factor DAF-7 in regulating *odr-10*. In larval hermaphrodites [40–42], *daf-7* expression in the ASI neurons decreases in response to nutritional stress. Previous work has shown that this signal is involved in many food-related aspects of *C. elegans* development and physiology, including entry into the dauer stage, fertility, fat metabolism, satiety quiescence, immune response, and lifespan [40–53]. *daf-7* also regulates the expression of several chemoreceptor genes in adult hermaphrodites, though *odr-10* was not found to be among these [54]. Interestingly, a recent study found that *daf-7* is male-specifically expressed in the ASJ neurons, where it promotes food-leaving behavior [38].

We found that well-fed *daf-7* mutant males displayed significant upregulation of *ODR-10::GFP* (Fig. 3A), indicating that this signal also has a role in repressing *odr-10* expression. This regulation appears to be male-specific, as we detected no change in *ODR-10::GFP* in *daf-7* mutant hermaphrodites (Fig. 3A). To further explore this, we genetically masculinized the nervous system of hermaphrodites by pan-neural expression of the male sex-determination regulator*fem-3* [19, 21]. Consistent with previous work [12], this was sufficient to reduce expression of *ODR-10::GFP* to male-typical levels (Fig. 3B). Further, pan-neural masculinization made hermaphrodite *odr-10* expression sensitive to *daf-7* loss, with *odr-10* levels increasing to an extent comparable to that seen in wild-type males (Figs. 3A, B). Thus, sex differences in the nervous system itself account for the male-specificity of the regulation of *odr-10* by *daf-7*. As pan-neural masculinization is sufficient to activate *daf-7* expression in ASJ in hermaphrodites [38], its expression in this neuron likely explains the sex-specificity of its role in regulating *odr-10*.

**Figure 3.**
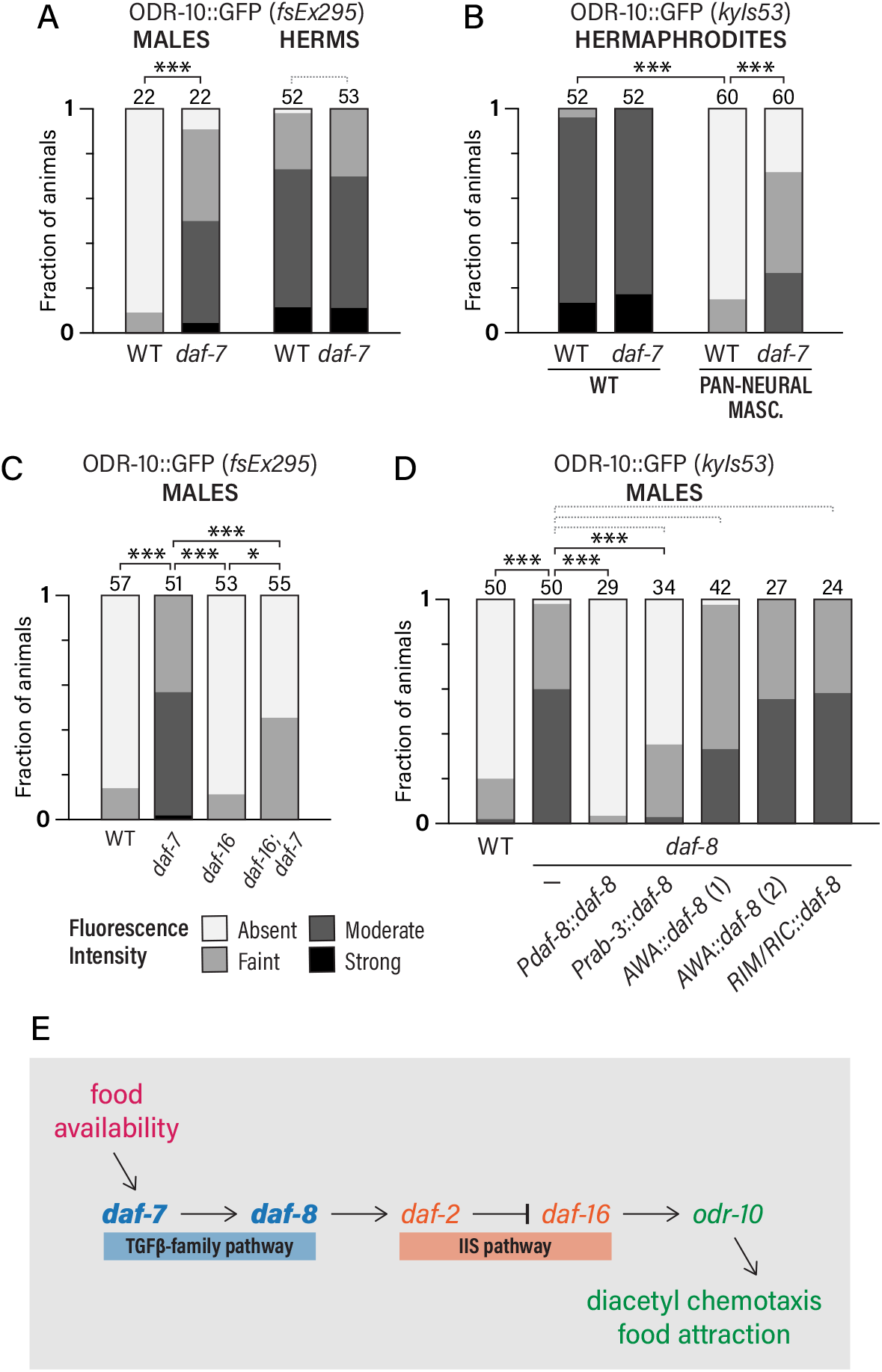
*odr-10* is sex-specifically regulated by *daf-7* TGFβ. (A) ODR-10::GFP expression in WT and *daf-7* males and hermaphrodites. (B) ODR-10::GFP expression in WT and *daf-7* hermaphrodites, with and without pan-neural masculinization by *Prab-3::fem-3(+)*. (C) ODR-10::GFP expression in WT, *daf-7, daf-16*, and *daf-16; daf-7* mutant males. (D) ODR-10::GFP expression in WT and *daf-8* mutants carrying the indicated *daf-8* expression constructs. (E) Pathway for the regulation of *odr-10* by *daf-7* and *daf-2* (IIS) signaling. *0.01 < *p* < 0.05; **0.001 < *p* < 0.01; \*\*\**p* < 0.001. Dotted gray brackets indicate *p* > 0.05.

Next, we explored the relationship between IIS and *daf-7* signaling. Crosstalk between these pathways has been found previously: several insulin signaling peptide genes are transcriptionally regulated by *daf-7* signaling [55, 56] and *daf-7* mutants show increased nuclear localization of the IIS effector DAF-16 [53, 57]. Consistent with this, we found that *daf-16* was required for the strong induction of *odr-10* in *daf-7* mutant males, such that *ODR-10::GFP* intensity was only slightly higher in *daf-16; daf-7* double mutants than in *daf-16* single mutants (Fig. 3C). This suggests that *daf-7* primarily regulates *odr-10* expression by modulating IIS, with secondary effects through other mechanisms (Fig. 3E).

To better understand the regulation of *odr-10* by *daf-7*, we sought to determine the cellular focus of TGFβ pathway activity. First, we found that males carrying a mutation in *daf-8*, a SMAD downstream of *daf-7* [58], phenocopied the high *odr-10* expression of *daf-7* mutants (Fig. 3D). Further, expression of a *daf-8* cDNA under the control of its own promoter or a pan-neural promoter was sufficient to reduce *odr-10* expression in *daf-8* mutant males (Fig. 3D).

However, we detected no rescue when we expressed *daf-8* specifically in AWA (Fig. 3D). Similarly, expression of *daf-8* in RIM and RIC, neurons known to be modulated by *daf-7* to control dauer formation, pharyngeal pumping, and fat accumulation [42], had no effect on *odr-10* expression (Fig. 3D), though it did rescue dauer formation (not shown). Thus, *daf-8* likely acts in the nervous system, but its activity in AWA, RIM, or RIC is not sufficient for the feeding-state regulation of *odr-10* in males.

### *Food perception, rather than internal metabolic state, regulates* odr-10 *in males*

During food deprivation, animals experience both metabolic stress and the loss of exposure to chemical signals from food itself. To explore the relative roles of these on *odr-10* expression, we used several methods to uncouple physiological state from the detection of food cues. First, we fed worms *E. coli* OP50, the standard laboratory diet, pre-treated with the cytokinesis inhibitor aztreonam. This generates long chains of bacteria that cannot be ingested by *C. elegans* [59]. Confirming this, we observed no accumulation of GFP inside nematodes cultured on aztreonam-treated GFP-labeled *E. coli*, even though GFP was clearly visible in the pharynx and intestine of animals exposed to control GFP-labeled bacteria (Fig. S3). Moreover, the intestine of animals cultured on this inedible food was clearly distended, similar to animals deprived entirely of bacterial food (Fig. S3). Despite this metabolic stress, however, these animals displayed only a very modest increase in *ODR-10::GFP* expression (Fig. 4A). Next, we re-exposed food-deprived animals to either control or aztreonam-treated bacteria for 24 h. In both cases, *ODR-10::GFP* expression returned to baseline levels, even though those cultured on aztreonam-treated bacteria presumably remained in a starved state (Fig. 4B). Thus, while metabolic stress may play a small role in *odr-10* activation upon food deprivation in adult males, the loss of chemosensory cues appears to be the primary driver of this regulation.

**Figure 4.**
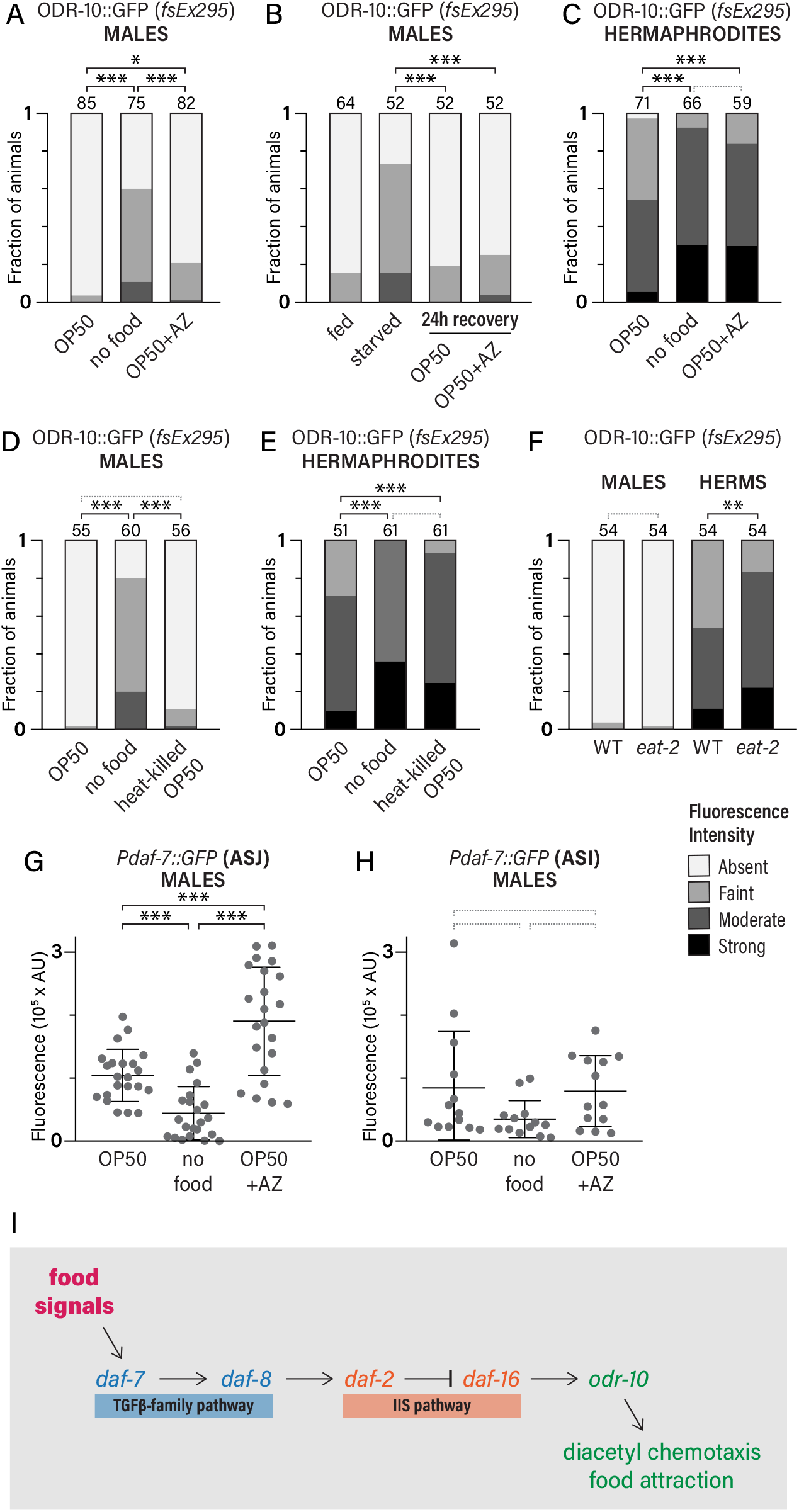
Sensory perception, not a metabolic cue, regulates *odr-10* and *daf-7* in males. (A) ODR-10::GFP expression in males cultured on *E. coli* OP50, without food, or on aztreonam-treated OP50 for 16-18 h. (B) ODR-10::GFP expression in well-fed males, food-deprived males, and food-deprived males after 24 h recovery on control or aztreonam-treated OP50. (C) ODR-10::GFP expression in hermaphrodites cultured on OP50, without food, or on aztreonam-treated OP50 for 16-18 h. (D-E) ODR-10::GFP expression in males (D) and hermaphrodites (E) cultured on OP50, without food, or on heat-killed OP50 for 16-18 h. (F) ODR-10::GFP expression in WT and *eat-2* males and hermaphrodites. (G-H) *Pdaf-7::GFP* fluorescence intensity in ASJ (G) and ASI (H) on males cultured on OP50, without food, or on aztreonam-treated OP50. (I) Pathway for the regulation of *odr-10* by TGFβ and IIS in response to external food signals in males. *0.01 < *p* < 0.05; **0.001 < *p* < 0.01; \*\*\**p* < 0.001. Dotted gray brackets indicate *p* > 0.05.

To ask whether this principle applies in both sexes, we exposed adult hermaphrodites to aztreonam-treated bacteria. Unlike males, hermaphrodites responded with full induction of *ODR-10::GFP*, mounting a response indistinguishable from that elicited by food deprivation (Fig. 4C). This indicates that the role of food signals in the regulation of *odr-10* differs markedly by sex, such that the induction of *odr-10* in food-deprived hermaphrodites appears to result primarily from nutritional stress, rather than the loss of chemosensory cues.

This apparent sex-specific sensitivity to nutritional status suggested that reducing food quality or intake might increase *odr-10* expression in hermaphrodites but not in males. To test this, we grew animals on a low-quality food source, heat-killed *E. coli*, for 16-18 h. Under these conditions, *odr-10* expression in males was comparable to that observed in males grown on control food, but expression in hermaphrodites was significantly increased (Figs. 4D, E). We also examined the effects of reducing food intake by assaying *ODR-10::GFP* in *eat-2* mutants, in which the pharyngeal pumping rate and the intake of food are reduced [60]. We found that this had no detectable effect on *odr-10* expression in males, but that *odr-10* expression in *eat-2* hermaphrodites was significantly higher than in control animals (Fig. 4F). These findings reinforce the idea that *odr-10* is regulated primarily by food-derived sensory signals in males and by internal metabolic state in hermaphrodites.

### tax-2/tax-4-*dependent sensory signals repress* odr-10 *expression via* daf-7 *and IIS*

Previous work has found that the male-specific expression of *daf-7* in ASJ depends on the well-fed state, but that expression in ASI does not [38]. To ask whether the effects of food signals on *odr-10* expression in males might be mediated by *daf-7* signaling, we examined *Pdaf-7::GFP* expression in animals cultured on aztreonam-treated *E. coli*. In control experiments, we verified that the male-specific expression of *Pdaf-7::GFP* in ASJ was dramatically reduced upon food deprivation, but that expression in ASI was relatively unchanged (Fig. 4G, H). In contrast, culturing animals on inedible food did not bring about downregulation of *daf-7* expression in ASJ or in ASI (Fig. 4G); on the contrary, for unknown reasons, *daf-7* expression in ASJ increased when males were grown on inedible food (Fig. 4G). This suggests that, in males, *daf-7* levels in ASJ are an internal representation of the detection of food rather than the level of physiological satiety. Moreover, these findings indicate that a reduction of *daf-7* expression couples decreased food perception to the activation of *odr-10* in adult *C. elegans* males.

If *odr-10* is regulated primarily by food detection in males, the disruption of sensory function should alter *odr-10* expression. To test this, we examined animals carrying mutations in *tax-2* and *tax-4*, genes that encode subunits of a cyclic nucleotide-gated cation channel required for the function of a subset of *C. elegans* chemosensory neurons [61, 62]. We found significant upregulation of *odr-10* in well-fed *tax-2* and *tax-4* mutant males (Fig. 5A), supporting this hypothesis. Notably, *tax-2* and *tax-4* are not expressed in AWA and are not required for AWA-mediated chemosensory behavior [63], indicating that sensory perception regulates *odr-10* non-cell-autonomously. In hermaphrodites, loss of *tax-2* caused a modest increase in *odr-10* expression, while loss of *tax-4* resulted in a trend toward increased expression (Fig. 5B). Thus, while chemosensory signals are not the primary modulator of *odr-10* expression in hermaphrodites, they likely contribute to its regulation.

**Figure 5.**
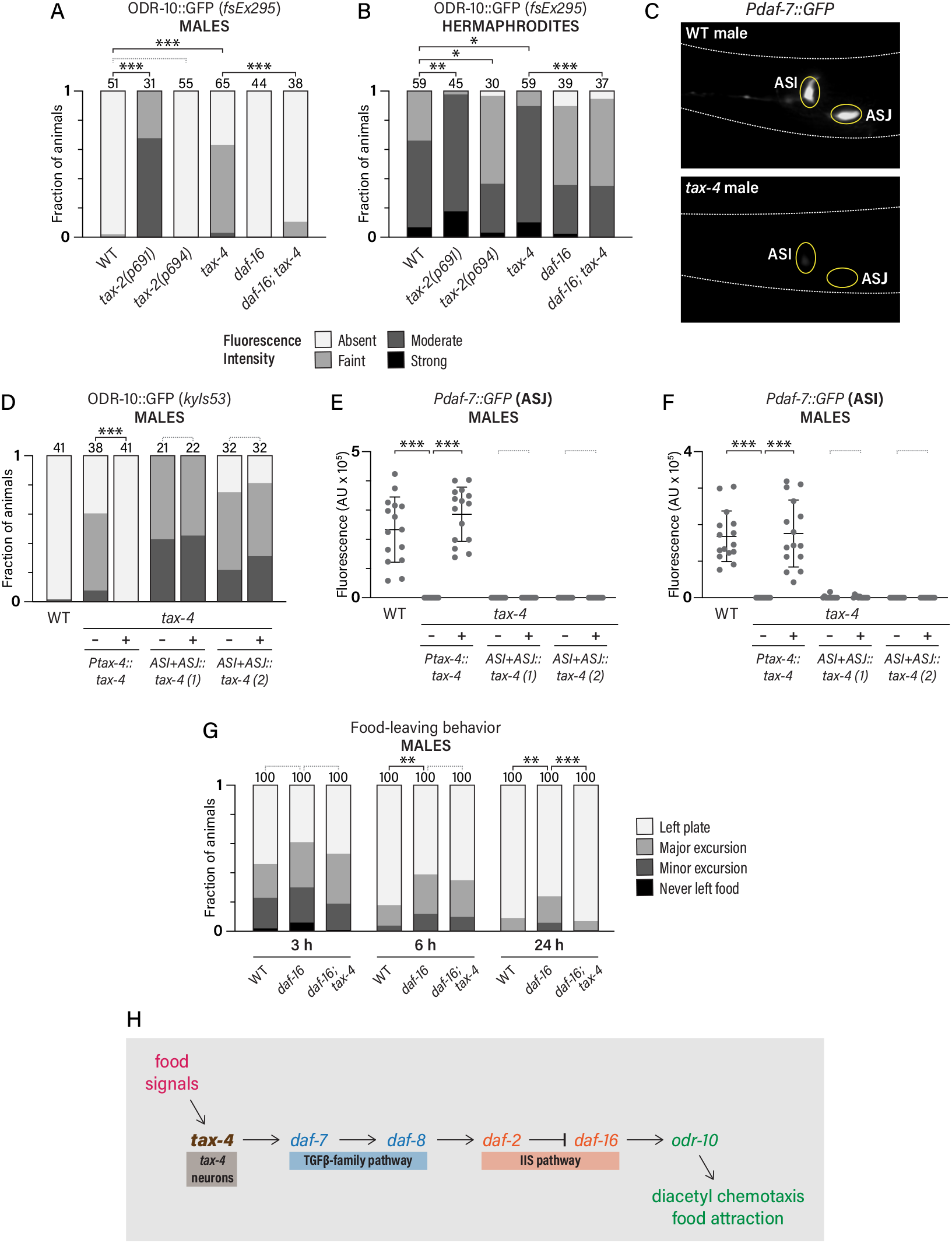
Signals from TAX-2/TAX-4 neurons regulate *odr-10* expression in males. (A-B) ODR-10::GFP expression in *tax-2, tax-4, daf-16*, and *daf-16; tax-4* males (A) and hermaphrodites (B). (C) Representative images of *Pdaf-7::GFP* expression in WT and *tax-4* males. Yellow ovals indicate the locations of ASI and ASJ. (D) ODR-10::GFP *(kyIs53)* expression in WT and *tax-4* males with the indicated *tax-4* expression constructs. Two separate lines are shown for ASI+ASJ rescue. (E-F) *Pdaf-7::GFP* fluorescence in ASJ (E) and ASI (F) in *tax-4* males with the indicated *tax-4* expression constructs. Two separate lines are shown for ASI+ASJ rescue. (G) Food-leaving behavior of wild-type, *tax-4*, and *daf-16; tax-4* mutant males at the indicated timepoints. For each plate, track patterns were scored manually using the four-point scale shown in the legend. (H) Pathway for the regulation of *odr-10* by food signals, TGFβ signaling, and IIS in males. *0.01 < *p* < 0.05; **0.001 < *p* < 0.01; *\*\*\*p* < 0.001. Dotted gray brackets indicate *p* > 0.05.

If sensory signals from *tax-4*-expressing neurons repress *odr-10* expression via *daf-7* and IIS, the elevated *odr-10* expression in *tax-4* mutant males should depend on *daf-16*. Indeed, we found that the *odr-10* expression phenotype of *tax-4* males was strongly suppressed in *daf-16; tax-4* double mutants (Fig. 5A). Similarly, *daf-16* was essential in hermaphrodites for the modest effect of *tax-4* loss on *odr-10* expression (Fig. 5B). Thus, the sensory signals mediated by *tax-4*-positive neurons likely act through IIS to repress *odr-10* expression. Interestingly, previous work has shown that the expression of *daf-7* is reduced in *tax-4* mutant hermaphrodites [45], suggesting that the increase in *odr-10* expression we observed in *tax-4* males could result from a reduction in *daf-7* expression in ASJ and/or ASI. Consistent with this, we found that *Pdaf-7::GFP* expression in *tax-4* males was strongly reduced in ASI and was virtually undetectable in ASJ (Fig. 5C, E, F). Thus, the high expression of *odr-10* in *tax-4* males likely results from a reduction in *daf-7* expression. Together, this suggests that removal from food decreases the activity of a subset of *tax-4* neurons, leading to a reduction in *daf-7* production in ASJ, reduced IIS pathway activity, and increased *odr-10* expression.

The TAX-2/TAX-4 channel is required for sensory transduction in roughly 12 classes of *C. elegans* sensory neurons, including ASI and ASJ [61, 62]. To ask whether *tax-4* acts cell-autonomously in these neurons to promote the expression of *daf-7*, we carried out cell-type-specific rescue experiments. Although rescue of *tax-4* under the endogenous *Ptax-4* promoter reduced the elevated *odr-10* expression of *tax-4* mutant males, we observed no such effect when *tax-4* was expressed specifically in ASJ, ASI, or both neurons simultaneously (Figs. 5D and S4A). Consistent with this, ASI/ASJ-specific expression of *tax-4* did not restore *daf-7* expression in *tax-4* mutants (Fig. 5E, F). These results suggest that sensory perception regulates *daf-7* non-cell-autonomously, and that chemosensation in ASI and ASJ does not regulate *odr-10*.

To narrow the focus of sensory function required for *daf-7* and *odr-10* expression, we used *tax-2(p694)*, an unusual allele that limits *tax-2* function to ASI, ASJ, and four other pairs of neurons: AWB, AWC, ASK, and ASG [61, 64]. Unlike *tax-2* null mutants, *tax-2(p694)* males showed no increase in *odr-10* expression (Fig. 5A), indicating that one or more of this group of neurons is sufficient to transduce the sensory signals that repress *odr-10* in males on food. Examining these classes individually, we found that *ceh-36* mutant males, in which AWC does not properly differentiate, displayed no changes in *odr-10* expression (Fig. S5A). Similarly, *odr-10* was unaltered in animals in which AWB or ASK was genetically ablated (Fig. S5B, C). Furthermore, we found that ASG-specific expression of *tax-4* failed to reduce *odr-10* expression in *tax-4* males (Fig. S5D). Thus, we conclude that, in well-fed males, *odr-10* expression is repressed by the distributed detection of food-derived chemosensory cues (Fig. 5G).

### tax-2/tax-4-*dependent sensory signals, acting through IIS, contribute to the balance between feeding and exploration*

Previously, we showed that overexpression of *odr-10* in adult males was sufficient to reduce food-leaving, while loss of *odr-10* in adult hermaphrodites increased it [12]. The size of these effects is relatively small, consistent with the idea that modulation of *odr-10* is just one of several mechanisms regulating this behavior [32]. Therefore, we sought to assess the contribution of the feedback loop we identify here to the feeding-vs.-exploration decision. To do so, we measured food-leaving behavior in wild-type, *tax-4*, and *daf-16; tax-4* males. *tax-4* mutant males should mimic a state of prolonged food absence, with low *daf-7* and high *odr-10* expression inhibiting food-leaving behavior [12, 38]. However, because TAX-2/TAX-4 neurons are themselves important for food detection [65], loss of *tax-4* function seems likely to increase food-leaving, making it difficult to predict how the behavior of these animals might differ from wild-type. We found that *tax-4* mutant males exhibited modestly reduced food-leaving compared to WT males (Fig. 5G), suggesting that the strong food-leaving defect expected to result from low *daf-7* expression [38] is counteracted by the absence of food detection by TAX-2/TAX-4 neurons.

Because *odr-10* activation in *tax-4* mutants requires *daf-16*, the loss of *daf-16* might be expected to increase the food-leaving behavior of *tax-4* males. Indeed, at 24 hr, *daf-16; tax-4* males exhibited slightly more food-leaving behavior than did *tax-4* males (Fig. 5G). Because constitutive loss of *tax-4* and *daf-16* likely causes broad changes in worm behavior and physiology, these results should be interpreted cautiously. Nevertheless, they are consistent with the idea that the sensory- and *daf-16*-dependent modulation of *odr-10* contributes to recalibrating the feeding-vs.-exploration balance in food-deprived males.

## DISCUSSION

Homeostatic regulation is a central feature of many animal behaviors and drives, including feeding, sleep, and reproduction, and is often driven by internal signals representing aspects of physiological and metabolic state [66]. In *C. elegans* males, *odr-10*, the chemoreceptor for the food-associated odor diacetyl, helps tune the behavioral equilibrium between mate-searching and feeding. In well-fed adults, low expression of *odr-10* reduces food attraction and promotes food-leaving; food deprivation upregulates *odr-10*, contributing to increased food attraction and decreased food-leaving [12]. Here, we identify a neuroendocrine feedback mechanism that regulates *odr-10* expression as a function of feeding state. Surprisingly, the low *odr-10* expression typical of well-fed adult males is not enabled by signals that reflect nutritional status. Rather, the presence of food itself engages a neuroendocrine feedback mechanism that downregulates *odr-10*, blunts chemosensory acuity, and helps balance the nutritional and reproductive needs of the adult *C. elegans* male.

In the model we propose (Fig. 6), multiple food-derived chemosensory cues are detected by a subset of sensory neurons that express a cyclic-nucleotide-gated channel comprising the subunits TAX-2 and TAX-4. Our results suggest that multiple TAX-2/TAX-4-positive neurons contribute at this step. Using an unusual allele of *tax-2*, we find that the activity of six pairs of chemosensory neurons (AWB, AWC, ASG, ASI, ASJ, and ASK) is sufficient to provide the “on-food” signal. Rescue and ablation experiments indicates that this signal is distributed among multiple members of this group, at least four of which are known to detect food-associated signals [65]. Because bacterial food sources emit complex and variable chemical signatures [67], it is not surprising that adult males use multiple streams of information to assess the presence of food.

**Figure 6.**
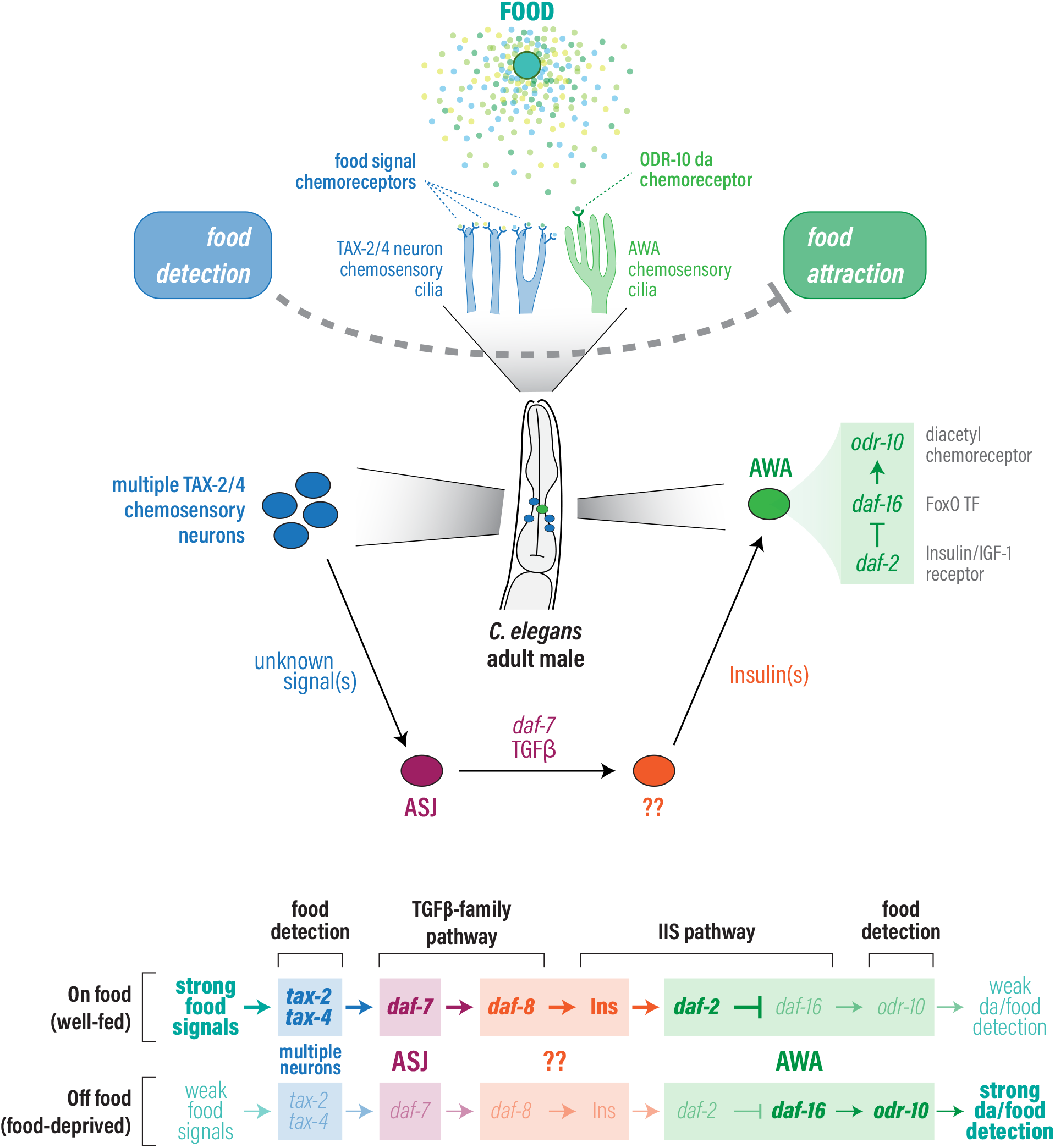
A sex-specific chemosensory feedback loop couples food detection, TGFβ signaling, and IIS to expression of the chemoreceptor *odr-10* in adult *C. elegans* males. The model shows the proposed mechanism by which chemosensory information about food availability is transmitted through a neuroendocrine loop to regulate *odr-10* in AWA. Pathways at the bottom depict the genetic architecture of the mechanism. See Discussion for details.

Next, these sensory signals converge to regulate expression of the TGFβ-superfamily ligand DAF-7. Production of this signal by the ASI neurons has been repeatedly shown to represent favorable environmental conditions, both in larvae and adults, and *daf-7* expression is known to be regulated by sensory signals in hermaphrodites [40–45, 48, 49, 51, 68]. Interestingly, recent work has identified a sex difference in *daf-7* expression, such that well-fed adult males also produce DAF-7 in the ASJ neurons, where its expression promotes food-leaving behavior [38]. Our results indicate that *daf-7* expression in ASJ responds to sensory information from TAX-2/TAX-4 neurons: both in the absence of food and in *tax-4* mutants, *daf-7* expression in ASJ is markedly reduced. Interestingly, although both ASJ and ASI are TAX-2/TAX-4 neurons, we find that the sensory function of these cells alone is insufficient to activate *daf-7* expression in the presence of food. Thus, non-cell-autonomous signals from AWB, AWC, ASK, and/or ASG are transmitted to ASJ and ASI, indicating that these neurons integrate multiple dimensions of sensory information into a single environmental-state quality signal.

*daf-7* is known to influence many aspects of *C. elegans* development, physiology, and behavior, including decisions about whether to remain on a food source [38, 47]. Many of these effects are mediated by regulation of the RIC and RIM interneurons via the SMAD *daf-8* and other members of the canonical TGFβ signaling pathway [42, 58]. However, our results indicate that, with respect to the regulation of *odr-10* in adult males, *daf-8* does not act in RIC/RIM, nor in AWA, the neuron in which *odr-10* is expressed. However, because the *odr-10* expression defect of *daf-8* males can be rescued by pan-neural expression of *daf-8*, the *daf-7* signal must act somewhere in the nervous system.

In addition to the *daf-7* signal, signaling through the IIS pathway is important for maintaining low *odr-10* expression in well-fed adult males. Males bearing mutations in the IIS receptor *daf-2* exhibit high *odr-10* expression even in the well-fed state; this phenotype is suppressed by loss of the FoxO family factor *daf-16*, the primary effector of *C. elegans* IIS [69]. Importantly, loss of *daf-16* prevents the upregulation of *odr-10* upon food deprivation, indicating that, in wild-type adult males, removal from food leads to downregulation of *daf-2* activity and the activation of *daf-16*. Furthermore, the high *odr-10* expression seen in *daf-7* and in *tax-4* mutant males largely depends on *daf-16* function, strongly suggesting that these components of the feedback mechanism act upstream of IIS. Though *daf-7* and IIS pathways act largely independently in the dauer development decision [70–72], *daf-7* can also act upstream of IIS [53, 55–57]. Thus, we favor a model in which *daf-8* acts in the nervous system to regulate the production and/or release of one or more insulin-like peptides (ILPs). Because the *C. elegans* genome encodes 40 ILPs [69], further studies are needed to identify the specific ligands involved in this mechanism and the site of their production.

In the last steps of the feedback mechanism, one or more ILPs act directly on the AWA chemosensory neurons. Cell-type-specific rescue experiments demonstrate that *daf-16* acts in AWA; moreover, we find that *odr-10* is a very likely to be direct transcriptional target of *daf-16*. While *daf-16* is best known for its role in activating regulators of stress response and reprogramming metabolism, our results, together with those of others [5, 7, 73] show that IIS can also directly regulate chemosensory function. This increase in *odr-10* expression, likely just one part of a suite of changes in AWA physiology, increases sensitivity to diacetyl and to bacterial food. Interestingly, AWA has recently been found to play a male-specific role in detecting volatile (non-ascaroside) pheromones released by hermaphrodites [74]. One intriguing question for future work will be to ask whether the mechanism we describe here also regulates the pheromone-detection function of AWA, perhaps blunting it in food-deprived males. Regardless, the coupling of sensory function to chemoreceptor expression has precedent in *C. elegans* [7, 54, 75]; our work shows that this mechanism is deployed by *C. elegans* as a component of sex- and food-dependent behavioral plasticity.

While reproductive drives are well appreciated as key motivators of animal behavior, the mechanisms underlying sex differences in such drives remain largely mysterious. Biological sex can also influence other drives and their relative prioritization—for example, sleep homeostasis in *Drosophila* is subject to regulation by male-specific sexual arousal [76, 77]—but the mechanistic interactions between sex and other internal states has received little attention. We find that the chemosensory feedback loop described here is largely sex-specific. While food deprivation boosts *odr-10* expression in both sexes, only in males is this primarily driven by sensory feedback, *daf-7*, and the IIS-mediated activation of *odr-10*. In hermaphrodites, these mechanisms have relatively little contribution to *odr-10* regulation. Instead, a physiological cue, mediated by unknown signals, is the primary influence on *odr-10* in food-deprived hermaphrodites. The basis for this sex difference lies within the nervous system itself, as pan-neural masculinization of the hermaphrodite nervous system is sufficient to confer sensitivity to *daf-7* on *odr-10* expression. The recent finding of male-specific expression of *daf-7* in ASJ likely underlies the male-specificity of the mechanism we describe here, as the expression of *daf-7* in ASJ is particularly sensitive to food [38] and to sensory signals (this work) compared to that in ASJ. Furthermore, pan-neural masculinization (but not masculinization of ASJ alone) activated *daf-7* expression in ASJ in hermaphrodites [38]. Together with our finding that multiple TAX-2/TAX-4-family chemosensory neurons are important for *daf-7* and *odr-10* regulation in males, this suggests that male-specific properties of multiple sensory neurons might alter the regulatory logic by which sensory signals are integrated by *daf-7*.

Why might males and hermaphrodites use different mechanisms to couple feeding status to *odr-10* expression? Differences in the ways that the sexes invest resources to optimize reproductive fitness may offer a clue. The production of sperm is relatively cheap; males may instead allocate resources to generating higher levels of motor activity in the service of mate-searching [78, 79]. In contrast, the production of oocytes is resource-intensive, requiring the continuous synthesis of many macromolecular components. As such, it is possible that relying on direct measurements of food consumption and physiological state provides hermaphrodites with a more reliable indicator of the pressure to find new sources of nutrition. Numerous signaling mechanisms—including *daf-7* signaling; IIS; metabotropic signaling by glutamate, monoamines, and neuropeptides; the kinase AMPK; and RICTOR/TORC2 signaling—have been implicated in linking feeding and nutritional uptake to metabolism and behavior in hermaphrodites, *e.g*. [42, 43, 46–48, 80–96]. Other than *daf-7* and IIS, one or more of these “gut-to-brain” signals seem likely candidates for regulating *odr-10* in hermaphrodites. Further, the functional significance of increased *odr-10* expression in a food-deprived hermaphrodite is unclear, as hermaphrodites are already strongly attracted to food, even in the well-fed state. Perhaps higher *odr-10* expression makes food detection even more efficient or helps override inhibitory signals that prevent the consumption of lower-quality food in well-fed animals.

Why males use sensory rather than metabolic cues to regulate *odr-10* is unclear. This would seemingly allow rapid assessment of environmental state, but behavioral plasticity in food-leaving behavior and *odr-10* expression occurs over hours, not seconds or minutes. It may be that the integration of multiple chemosensory streams by *daf-7* occurs over a longer time scale, generating a time-averaged signal that reflects recently encountered environmental conditions. Further, the time needed for signaling downstream of *daf-7* also likely contributes to the delay between removal from food and the increase in ODR-10 abundance. Another issue worth considering is that males also use chemosensory information about the presence of food when deciding whether to copulate with a hermaphrodite. In the absence of food, males mate poorly with hermaphrodites, possibly to avoid investing in progeny that will be born into an unfavorable environment [97]. Interestingly, inedible food provides an environment nearly as permissive for mating success as does control food, indicating that males use chemosensory cues about food availability when making this decision [98]. The relationship of these cues to those that regulate *odr-10* are unknown, but it seems likely that they are related.

The importance of chemosensory cues in calibrating *odr-10* expression and, indirectly, the feeding-vs.-exploration balance, contributes to a growing appreciation of the importance of chemosensory signals in regulating behavior, metabolism, and internal state in *C. elegans*. Chemosensory cues have long been known to be important for the developmental decision to enter the dauer state, a long-lived, stress-resistant larval stage whose entry can be triggered by the absence of food and exposure to population-density pheromones, and the *daf-7* TGFβ and *daf-2* IIS pathways are the primary regulators of this decision [72]. In adult hermaphrodites, chemosensory cues themselves have been shown or suggested to alter gene expression, physiology, behavior, and lifespan *e.g*. [7, 42, 44, 46, 48, 68, 89, 99–102]. By establishing an important role for such signals in the male-specific regulation of chemosensory function, our work shows that chemosensory signals contribute to the balance between feeding and mating drives, and that they do so at least in part by influencing chemosensory function itself.

## ACKNOWLEDGMENTS

For discussion and critical feedback, we thank current and past members of the Portman lab and the Western New York Worm Group. We are particularly grateful to Deborah Ryan, who contributed the data shown in Figure S1B. Some strains used in this work were provided by the *Caenorhabditis* Genetics Center, which is funded by NIH Office of Research Infrastructure Programs (P40 0D010440). This work was supported by grants from the NIH (R01 GM108885 and R01 GM130136) to D.S.P.

## AUTHOR CONTRIBUTIONS

L.W. and D.P. designed the experiments; L.W. and R.M. conducted the experiments; L.W., R.M., and D.P. analyzed and interpreted the results; and L.W. and D.P. wrote the manuscript.

## EXPERIMENTAL PROCEDURES

### Strains

Strains were grown on *E. coli* OP50 using standard methods [103] and maintained at 20°C. Unless otherwise stated, all strains contained *him-5(e1490)* in order to obtain a high number of males; this is considered the wild-type for the purposes of these studies. Strains carrying mutations in *daf-2, daf-7*, or *daf-8* were maintained at 15°C to prevent dauer entry. In these cases, all paired control strains were also maintained at 15°C. Animals were transferred to 25°C (for *daf-2* strains) or 20°C (for *daf-7* and *daf-8* strains) as L4 larvae; scoring of one-day adults was carried out the following day. See Table S1 for a complete list of *C. elegans* strains used in this work.

### Quantification of *ODR-10::GFP, Podr-10::GFP*, and *Pdaf-7::GFP*

All animals used for experiments were sex-segregated as L4 larvae the day before scoring. One-day old adults were mounted on a 4% agarose pad and immobilized with levamisole. GFP fluorescence was observed using a 63x PlanApo objective on a Zeiss Axioplan 2. Due to the complexity of the structure of the AWA cilia, accurate computer-based quantification of ODR-10::GFP fluorescence intensity was not possible. ODR-10::GFP intensity was therefore scored using a scale of 0-3 (0=absent, 1=faint, 2=moderate, 3=bright) as previously described [12, 37]. When possible, the experimenter was blinded to genotype. For quantification of *Podr-10::GFP* and *Pdaf-7::GFP* expression, the integrated density of GFP was calculated using FIJI [104]. Images were obtained under constant imaging conditions with the cell of interest in center of the field. Background was calculated by taking the average mean fluorescence of three random sections of background within the area of the animal. Total cell fluorescence was calculated as (integrated density of GFP in the region of interest) – (mean background fluorescence x area of the region of interest).

### RNA isolation and Quantitative RT-PCR

To isolate RNA, one-day old adult hermaphrodites were collected and either fed or starved for 18 hours and then flash frozen. After cuticle disruption, total RNA was isolated using the RNeasy plus Mini kit (Qiagen) and samples were treated with DNase I (Invitrogen). cDNA was generated with the SuperScript III First-Strand Synthesis System (Invitrogen) using oligo(dT). Quantitative RT-PCR was performed on a Bio-Rad MyiQ2 icycler with iQ SYBR Green Supermix (BioRad) using 2 μl of cDNA with reactions in triplicate. Cycling conditions were: 1 cycle of 95°C for 5 min; 45 cycles of 95°C for 30 s, 60°C for 30 s, and 72°C for 45 s, followed by melting curve analysis to verify product specificity. Standard curves were generated for each primer using three serial dilutions of cDNA to determine an efficiency estimate for each run. Threshold cycles (Ct) were determined by the Bio-Rad iQ5 optical system software, which were used with efficiency estimates to calculate relative odr-10 gene expression by the Pfaffl analysis method [105]. Two reference genes, *cdc-42* and Y45F10D.4, were used for normalization [106]. *odr-10* expression in starved hermaphrodites is shown normalized to levels detected in well-fed hermaphrodites.

### Behavioral Assays

Diacetyl attraction assays and food-leaving assays were performed as described [12, 21, 107]. In all cases, the investigator was blinded to genotype.

### Aztreonam-treated OP50

*E. coli* OP50 was treated with aztreonam as described [97]. The day before experiments, *E. coli* OP50 was grown in LB medium at 37°C with shaking to log-phase growth. Aztreonam was then added to a final concentration of 10 μg/ml and cultures were incubated for 3 additional hours without shaking. Bacteria were then seeded onto freshly made (same-day) NGM plates containing aztreonam (10 μg/ml). To avoid transferring untreated bacteria, worms were washed at least 3 times with M9 buffer before being placed on aztreonam plates. Animals were scored after 16-18 h. For paired control experiments, starved animals were plated on unseeded NGM plates containing 10ug/ml aztreonam to control for any effects of aztreonam on gene expression.

### Molecular biology and generation of transgenic strains

All cDNAs were amplified from total RNA extracted from *him-5(e1490)* cultures. (See Table S2 for primer sequences.) The resulting cDNAs were cloned into pDONR221 and recombined via Multisite Gateway Cloning (Invitrogen). The Podr-10^ΔDAF-16^::GFP transgene was generated by Gibson assembly [108]. Plasmids were injected into animals at a concentration of 20-50 ng/ul of rescue construct and 100 ng/ul of Pelt-2::GFP co-injection marker.

### Starvation experiments

One-day old adults were washed at least three times with M9 buffer and then transferred to NGM plates with either *E. coli* OP50 or no food. For GFP quantification experiments, animals were scored after 16-18 h. For behavioral assays, animals were tested after 12 h.

### Heat killed *E. coli* OP50

Liquid cultures of *E. coli* OP50 were incubated at 75°C for 1 h. Heat-killed bacteria were then plated and used for experiments on the following day.

### Statistical analyses

For categorical data (*e.g*., fluorescence intensity of ODR-10::GFP and behavior in the food-leaving assay), data were analyzed using non-parametric tests. For pairwise comparisons, Mann-Whitney tests were carried out. For multiple comparisons, we used Kruskal-Wallis analysis with Dunn’s correction. For other analyses (*e.g*., of diacetyl chemotaxis and quantitated GFP fluorescence), data were assumed to be normally distributed. In these cases, Welch’s *t* tests were used for pairwise comparisons. For multiple comparisons, we used Brown-Forsyth and Welch one-way ANOVA tests with Holm-Sidak’s correction. In all experiments, only those comparisons necessary to test the hypotheses under consideration were carried out. For this reason, graphs do not indicate the results of all possible comparisons. Instead, all graphs show the results of all comparisons made. Dotted gray brackets indicate comparisons in which the *p* value was greater than 0.05. In cases where *p* was 0.05 or less, asterisks indicate *p* value ranges as follows: *0.01 < *p* < 0.05; **0.001 < *p* < 0.01; ***p < 0.001.

**TABLE S1.**
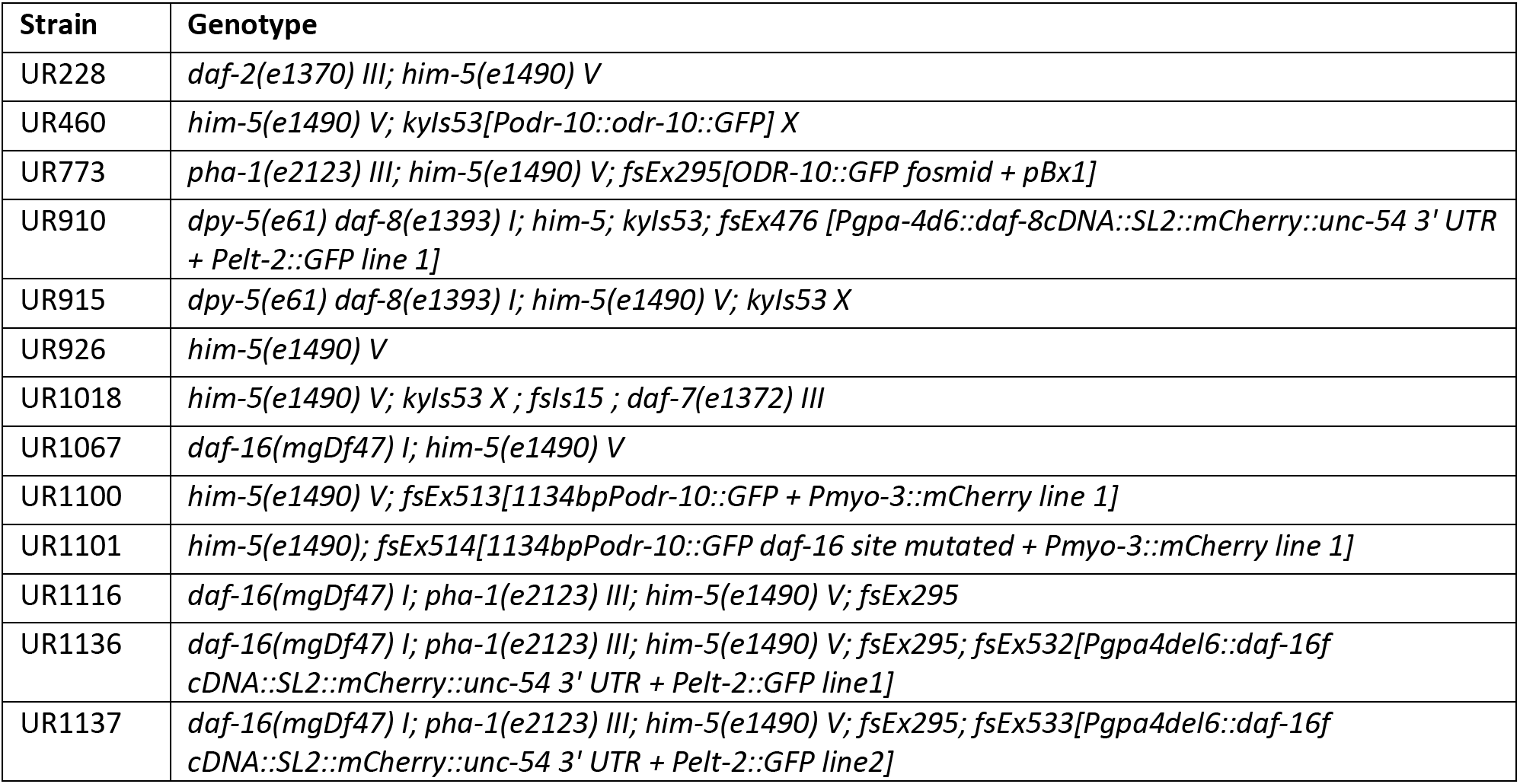
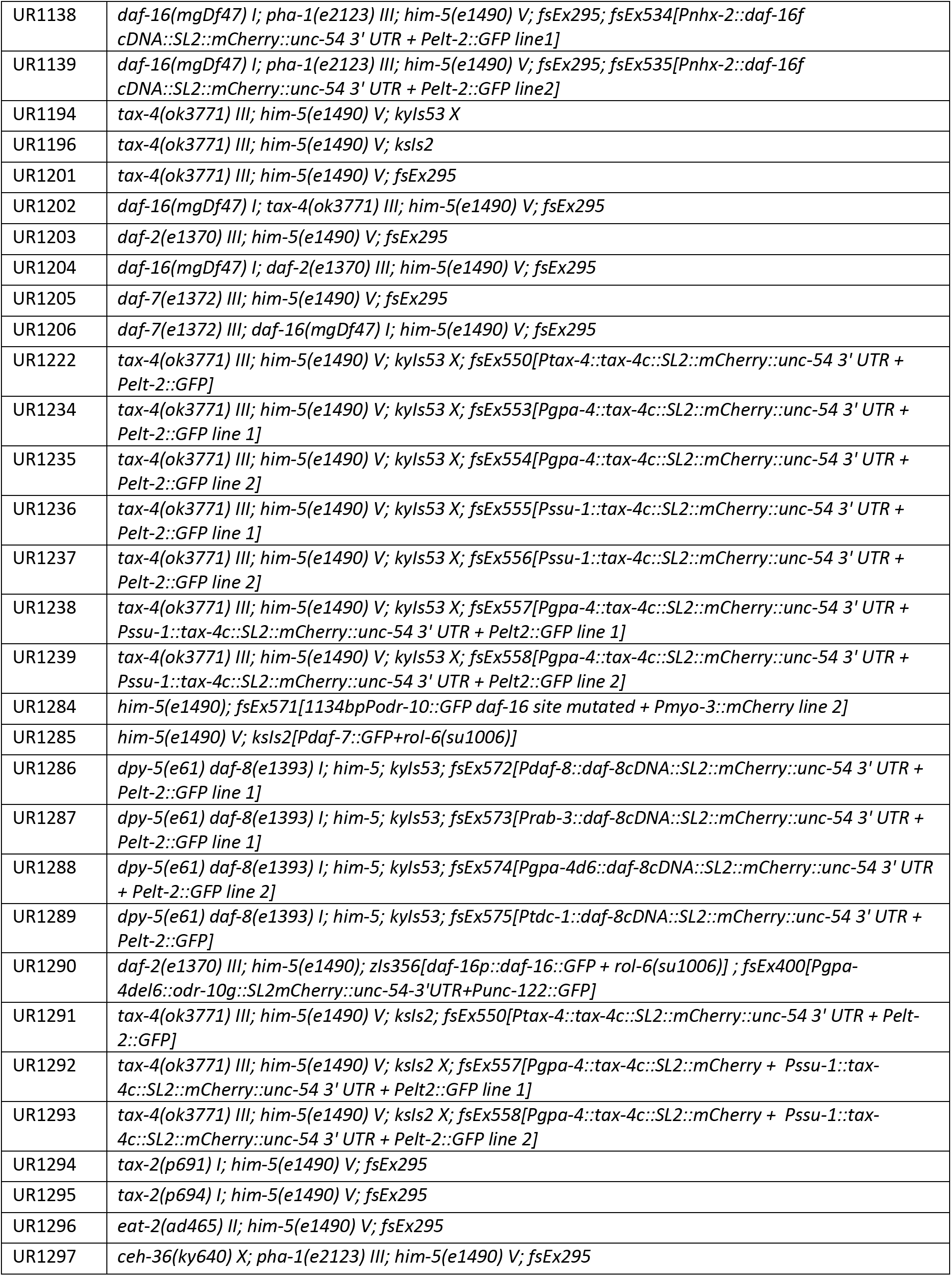
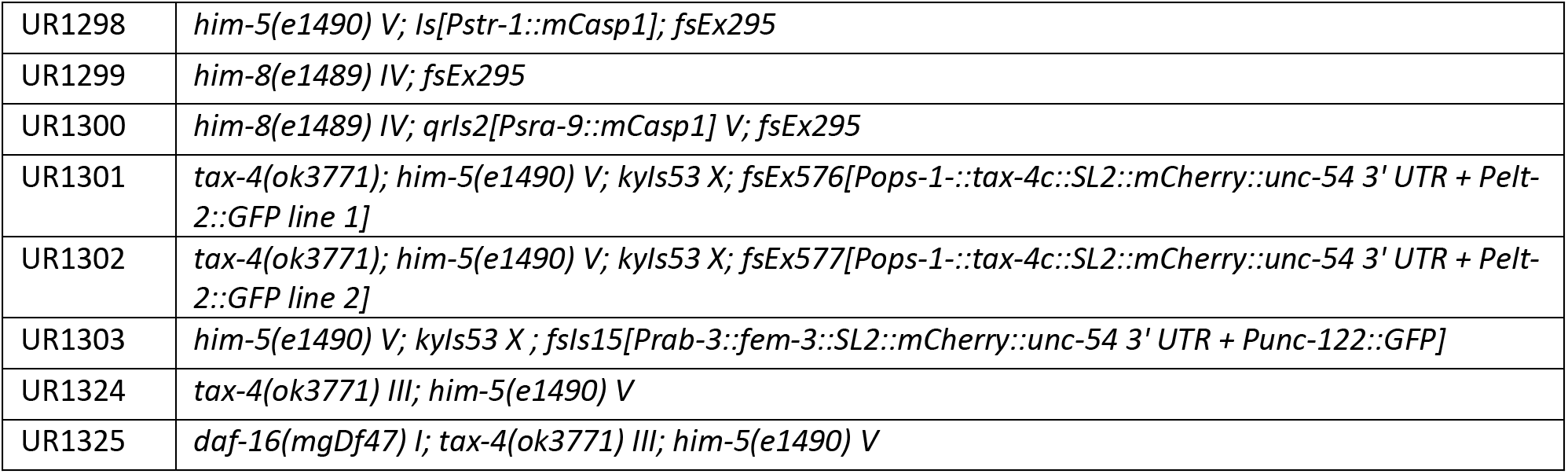
Strain List.

**TABLE S2.**
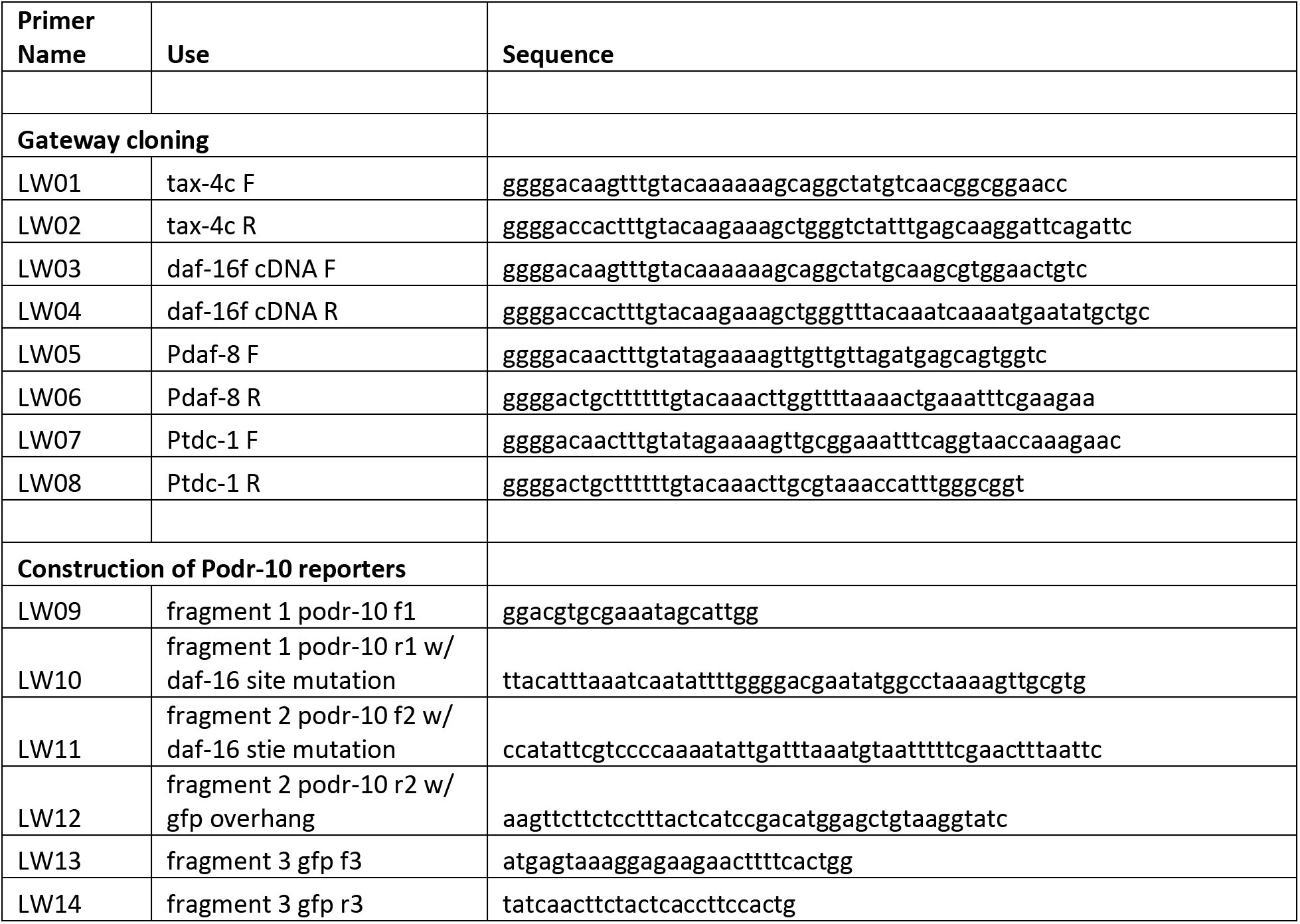
Primer List.

---

**Figure S1.**
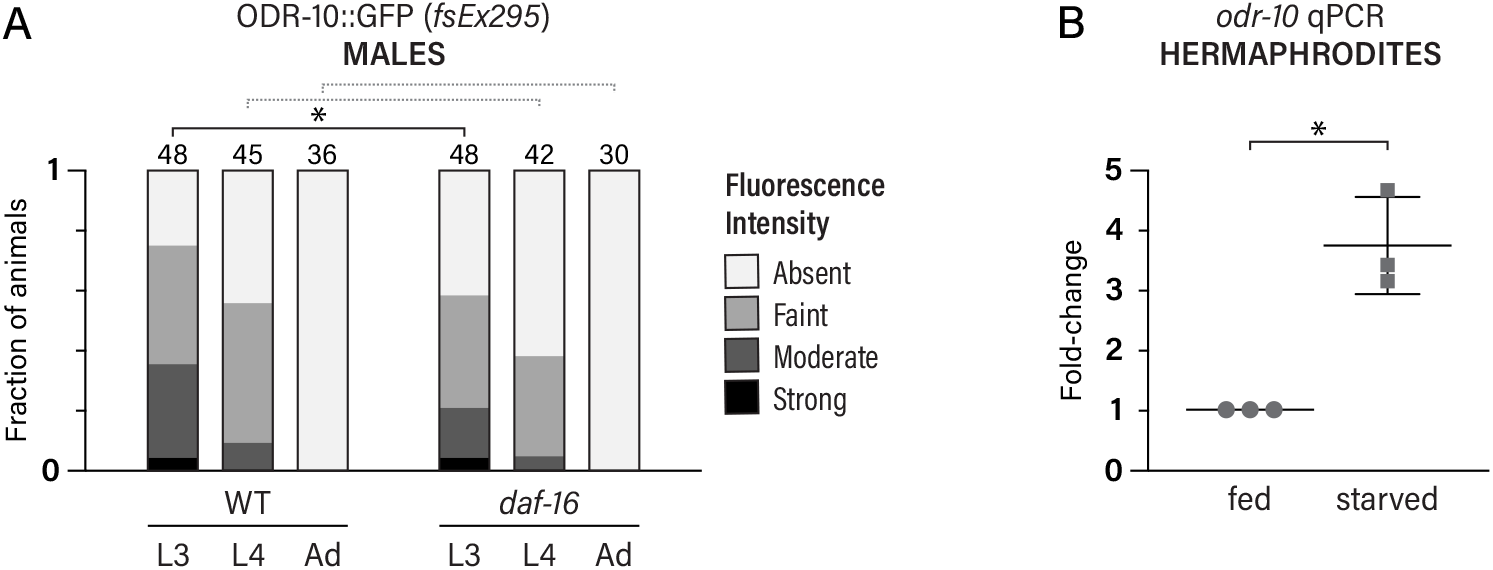
Related to Fig. 1. (A) ODR-10::GFP expression in L3, L4, and young adult WT and *daf-16* males. (B) *odr-10* mRNA levels as measured by qRT-PCR in fed and starved hermaphrodites, normalized to fed levels. *0.01 < *p* < 0.05; **0.001 < *p* < 0.01; \*\*\**p* < 0.001. Dotted gray brackets indicate *p* > 0.05.

**Figure S2.**
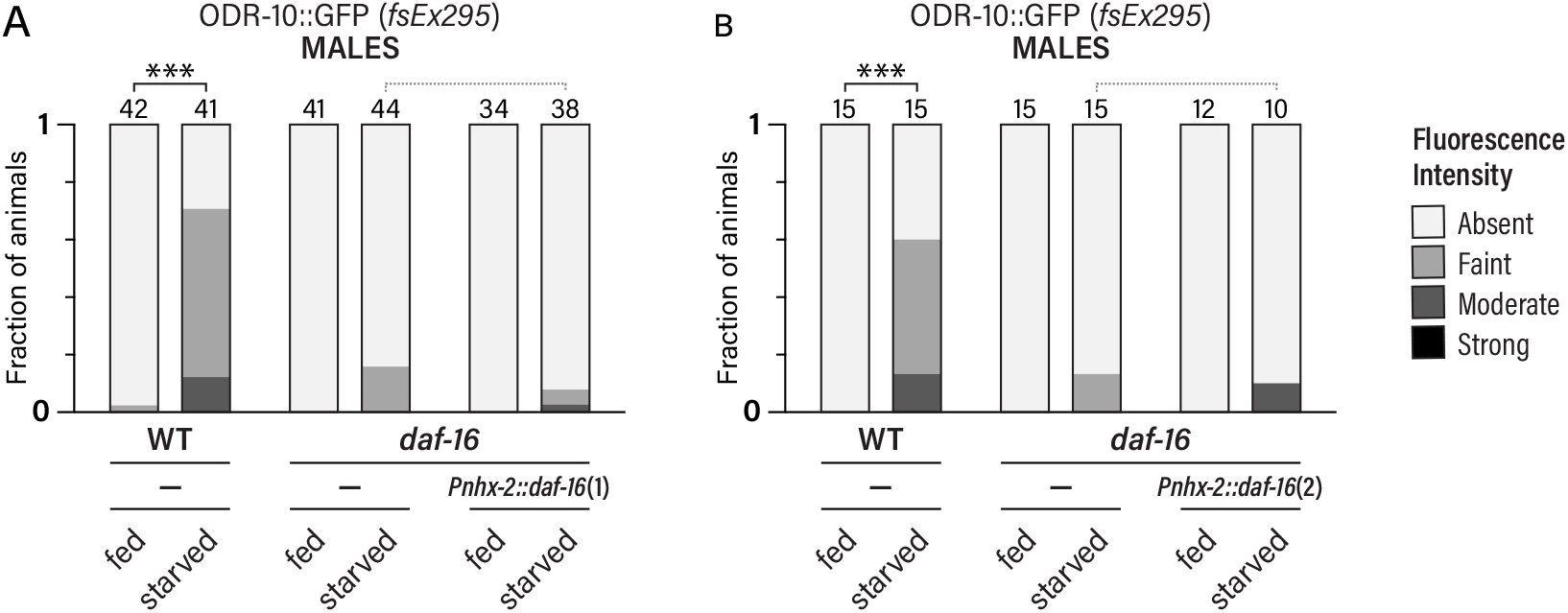
Related to Fig. 2. (A-B) ODR-10::GFP expression in WT and *daf-16* fed and starved males with or without transgenes expressing *daf-16f* cDNA in the intestine. Independent experiments for two independent transgenic lines are shown: (A) line 1; (B) line 2. *0.01 < *p* < 0.05; **0.001 < *p* < 0.01; \*\*\**p* < 0.001. Dotted gray brackets indicate *p* > 0.05.

**Figure S3.**
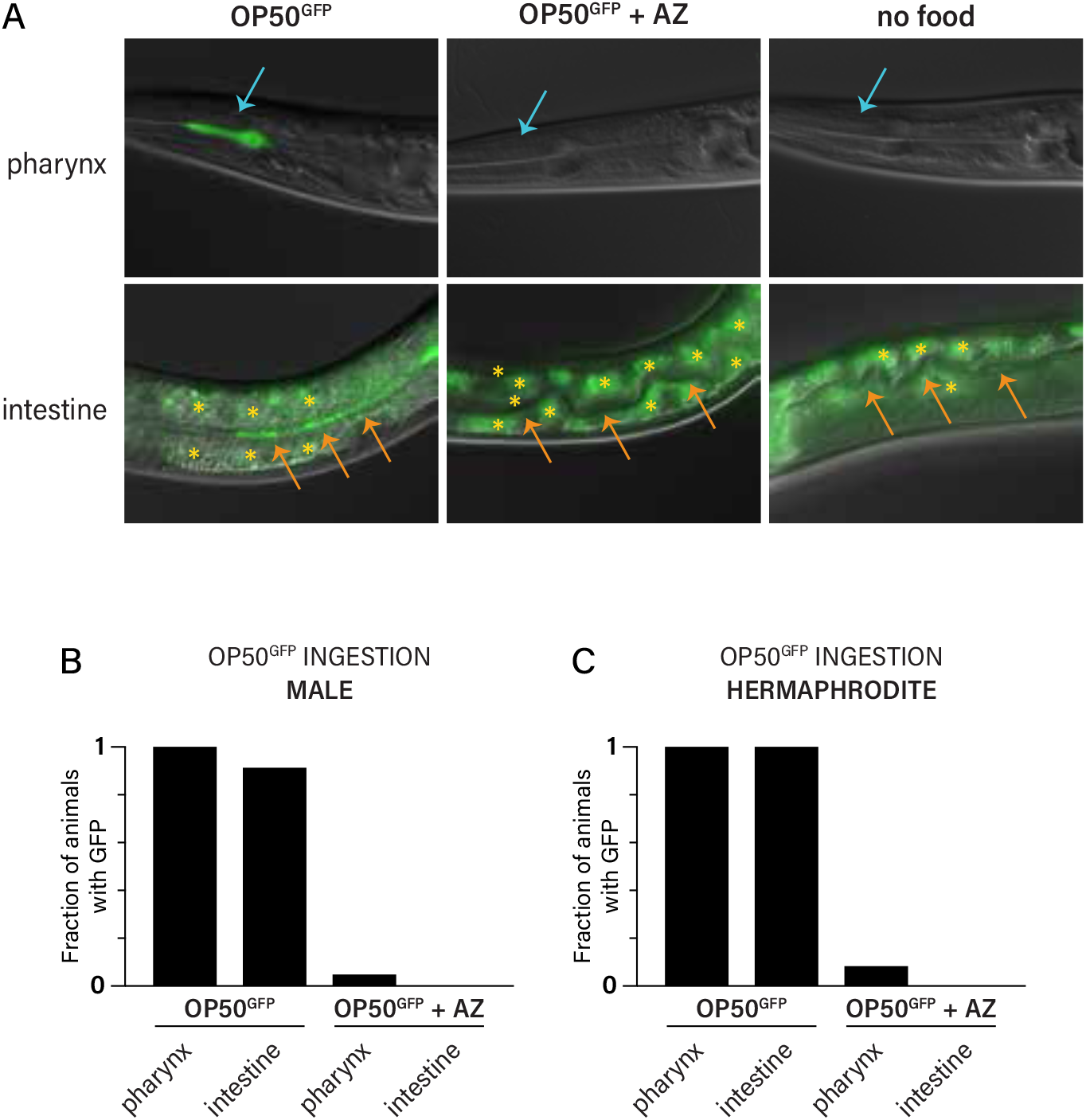
Related to Fig. 3. (A) Representative images of animals cultured with GFP-labeled *E. coli* OP50 (OP50^GFP^), with aztreonam-treated OP50^GFP^, or without food for 16-18 h. Blue arrows show the pharyngeal lumen; orange arrows indicate intestinal lumen. Asterisks indicate autofluorescent granules in the intestine. (B-C) Percent of animals with GFP detectable in the pharynx and intestine of animals fed OP50^GFP^ or aztreonam-treated OP50^GFP^ for 16-18 h.

**Figure S4.**
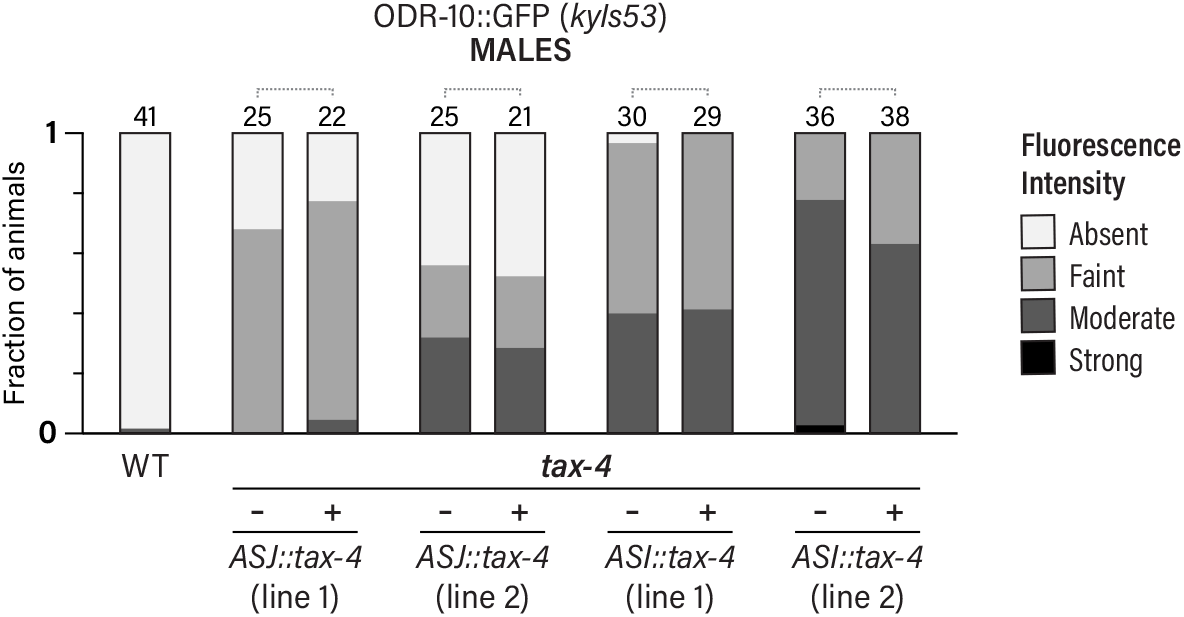
Related to Fig. 4. ODR-10::GFP (*kyIs53*) expression in WT and *tax-4* males with the indicated *tax-4* expression constructs. Dotted gray brackets indicate *p* > 0.05.

**Figure S5.**
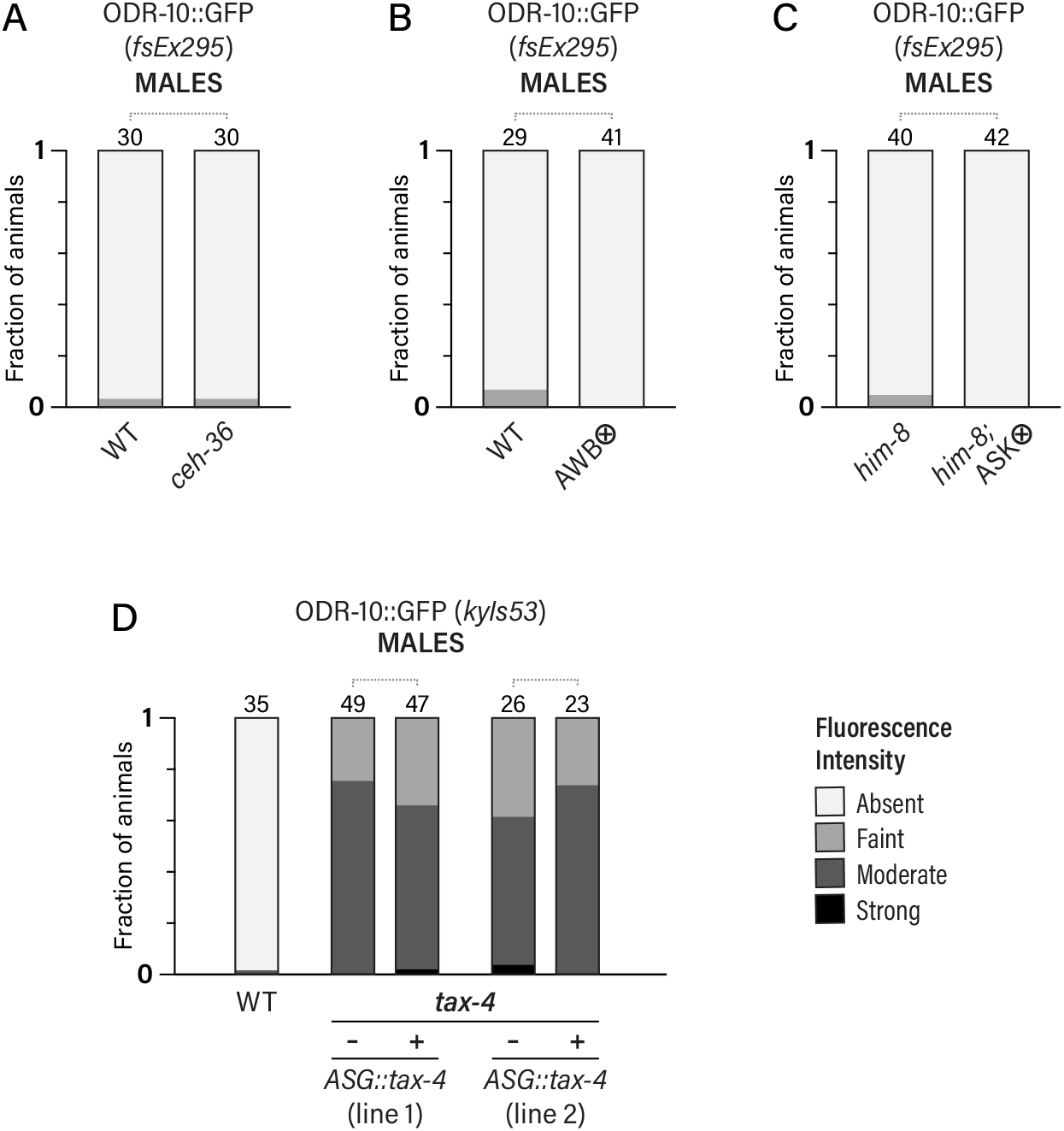
Related to Fig. 5. (A) ODR-10::GFP expression in WT and *ceh-36* males (AWC genetic ablation). (B) ODR-10::GFP expression in WT and *Pstr-1::mCasp1* males (AWB genetic ablation). (C) ODR-10::GFP expression in *him-8* and *him-8; Psra-9::mCasp1* males (ASK genetic ablation). (D) ODR-10::GFP (*kyIs53*) expression in WT and *tax-4* males with or without transgenes expressing *tax-4* cDNA in ASG (*Pops-1::tax-4c*). Results for two independent lines are shown. Dotted gray brackets indicate *p* > 0.05.

